# Presynaptic MAST Kinase Controls Bidirectional Post-Synaptic Responses to Convey Stimulus Valence in *C. elegans*

**DOI:** 10.1101/609479

**Authors:** Shunji Nakano, Muneki Ikeda, Yuki Tsukada, Xianfeng Fei, Takamasa Suzuki, Rhea Ahluwalia, Ayana Sano, Rumi Kondo, Kunio Ihara, Koichi Hashimoto, Tetsuya Higashiyama, Ikue Mori

## Abstract

Presynaptic plasticity is known to modulate the strength of synaptic transmission. However, it remains unknown whether regulation in presynaptic neurons alters the directionality –positive or negative-of postsynaptic responses. We report here that the *C. elegans* homologs of MAST kinase, Stomatin and Diacylglycerol kinase act in a thermosensory neuron to elicit in its postsynaptic neuron an excitatory or inhibitory response that correlates with the valence of thermal stimuli. By monitoring neural activity of the valence-coding interneuron in freely behaving animals, we show that the alteration between excitatory and inhibitory responses of the interneuron is mediated by controlling the balance of two opposing signals released from the presynaptic neuron. These alternative transmissions further generate opposing behavioral outputs necessary for the navigation on thermal gradients. Our findings reveal the previously unrecognized capability of presynaptic regulation to evoke bidirectional postsynaptic responses and suggest a molecular mechanism of determining stimulus valence.

## Introduction

Sensory stimuli that predict the valence of reward or punishment are major drivers of animal behaviors. For example, odors associated with predators are detrimental to the animals and elicit fear responses^1^, while smells predicting the presence of food or potential mates evoke different behaviors such as feeding^2^ or mating^3^. Extracting the valence information of sensory stimulus – whether the stimulus is attractive or aversive- and manifesting appropriate behavioral responses are the most vital function of the nervous system. Deciphering the molecular and circuit mechanisms underlying the valence coding of sensory information is thus fundamental to understand the principles of how animal behaviors emerge from the nervous system.

The valences of innate odor responses are guided by intrinsic property of the responding neurons, in which neurons expressing specific odorant receptors are hardwired in a neural circuit that elicits attractive or aversive behavior^4,5^. A similar labeled-line circuit operation was also demonstrated for encoding and responding to tastes such as sweet and bitter^6–8^, wherein cells expressing specific taste receptors are embedded in a specialized neural circuit that promotes or inhibits feeding behaviors.

Contrary to these developmentally programmed, stereotyped behaviors, the valence associated with certain sensory stimuli can vary depending on the past experience, the current environmental context and the stimulus intensity^9^. For example, olfactory preferences to the same odorants can differ depending on the odorant concentration^10^. Studies of worms, flies and mammals suggested a common feature of neural mechanism underlying the change in these odorant valences, wherein different concentrations of the same odorants recruit distinct sets of olfactory neurons and consequently change the perception of the same odorants^11–13^. However, the extent to which the brain utilizes different encoding strategies for alternating stimulus valence remains largely unexplored. In particular, the molecular and circuit mechanisms underlying the perception of altering valence for other sensory modalities are not yet understood.

The compact nervous system of *C. elegans* consisting of only 302 neurons provides an excellent opportunity to explore these questions^14^. *C. elegans* exhibits thermotaxis behavior^15^, in which the valence of thermal information varies depending on the past experience, current temperature environment and feeding states. Specifically, the temperature preference of *C. elegans* is plastic and determined by the cultivation temperature, in which animals that are cultivated at a constant temperature with food migrate toward that cultivation temperature on a thermal gradient without food^15^. When animals were placed at the temperature below the cultivation temperature they migrate up the thermal gradient, while above the cultivation temperature they move down the gradient, indicating that the valence associated with thermal stimuli alternates in opposing manners depending on the current temperature.

Previous studies identified neurons involved in thermotaxis^16–18^. Of those neurons, the AFD thermosensory neurons are essential for thermotaxis and are required for migrating up and down the thermal gradient to reach the cultivation temperature^17,19^. Calcium imaging analyses revealed that the AFD neuron displayed increases in calcium concentration upon warming phases of a temperature ramp both below and above the cultivation temperature^20,21^. However, it remains to be determined how the AFD neuronal activities with similar calcium dynamics below and above the cultivation temperature are transformed to encode appropriate valence of temperature information and the consequent manifestation of opposing behavioral regulations.

Here we report that the AFD neuron evokes opposing neuronal responses in its postsynaptic partner AIY interneuron below or above the cultivation temperature: the activation of AFD neuron stimulates the AIY neuron below the cultivation temperature, while it inhibits AIY above that temperature. We identified molecular components important for this process and showed that this alteration of the AFD-AIY communication is regulated by presynaptic actions of the *C. elegans* homologs of MAST (Microtubule-Associated Serine-Threonine) kinase^22^, Stomatin^23^ and Diacylglycerol kinase^24,25^. Our results further suggest that the alteration of the AFD-AIY synaptic transmission is mediated by the balance of two opposing signals released from the AFD neuron, an excitatory peptidergic signaling and an inhibitory glutamatergic signaling. A high-throughput behavioral analysis revealed that these alternative modes of the AFD-AIY transmission generate opposing behavioral biases in the steering direction of animal locomotion to warmer or colder side of the temperature gradient, thereby driving the animals toward the cultivation temperature. Our results suggest that bidirectional responses in the valence-coding neurons are regulated by presynaptic mechanism whereby the evolutionarily conserved MAST kinase, Stomatin and Diacylglycerol kinase control the presynaptic release of opposing signaling molecules.

## Results

### *kin-4, mec-2* and *dgk-1* Regulate the *C. elegans* Thermotaxis Behavior

To elucidate the molecular and neural mechanisms underlying the valence coding of thermal stimuli during *C. elegans* thermotaxis behavior, we conducted forward genetic screens and sought mutants that displayed abnormal thermotaxis behavior. We found that mutations in three genes, *kin-4, dgk-1* and *mec-2*, affected this behavior (Fig. 1). While the wild-type animals that had been cultivated at 20 °C preferred to stay around the cultivation temperature, loss-of-function mutants of *kin-4*, which encodes the *C. elegans* ortholog of the mammalian MAST (Microtubule Associated Serine Threonine) kinase, displayed a cryophilic phenotype and migrated toward a colder temperature region than the wild-type animals (Fig. 1b and Supplementary Fig. 1a). This defect was rescued by introduction of a *kin-4* genomic clone (Fig. 1c), indicating that *kin-4* is required for thermotaxis. Animals carrying mutations in the gene *dgk-1*, which encodes a homolog of Diacylglycerol kinase θ^24,25^, also preferred to migrate toward a colder temperature region, as was previously reported ^26^ (Fig. 1b and Supplementary Fig. 1a). The cryophilic phenotype of *dgk-1(nj274)* mutants was rescued by introduction of a *dgk-1* genomic clone (Supplementary Fig. 1b), indicating that *dgk-1* is required for thermotaxis. We also isolated a mutation in *mec-2*, which encodes a *C. elegans* homolog of Stomatin^23^. *mec-2(nj89)* animals harbored a mutation that is predicted to alter the glutamic acid 270 to a lysine (*E270K*, see Supplementary Fig. 1a), and displayed a thermophilic phenotype (Fig. 1b). Introduction of a mutant *mec-2(E270K)* clone into the wild-type animals phenocopied *mec-2(nj89)* mutants, while introduction of a wild-type *mec-2* clone into *mec-2(nj89)* mutants did not rescue the thermophilic defect (Fig. 1d). We also generated a null allele of *mec-2(nj251 Δ)* (Supplementary Fig. 1a) and found that *mec-2* null mutants were grossly normal in thermotaxis (Supplementary Fig. 1c), suggesting that some of the nine additional Stomatin genes encoded by the *C. elegans* genome could compensate the deficit of *mec-2*. These results indicate that *mec-2(nj89)* is a gain-of-function mutation and causes a thermophilic phenotype.

**Figure 1.**
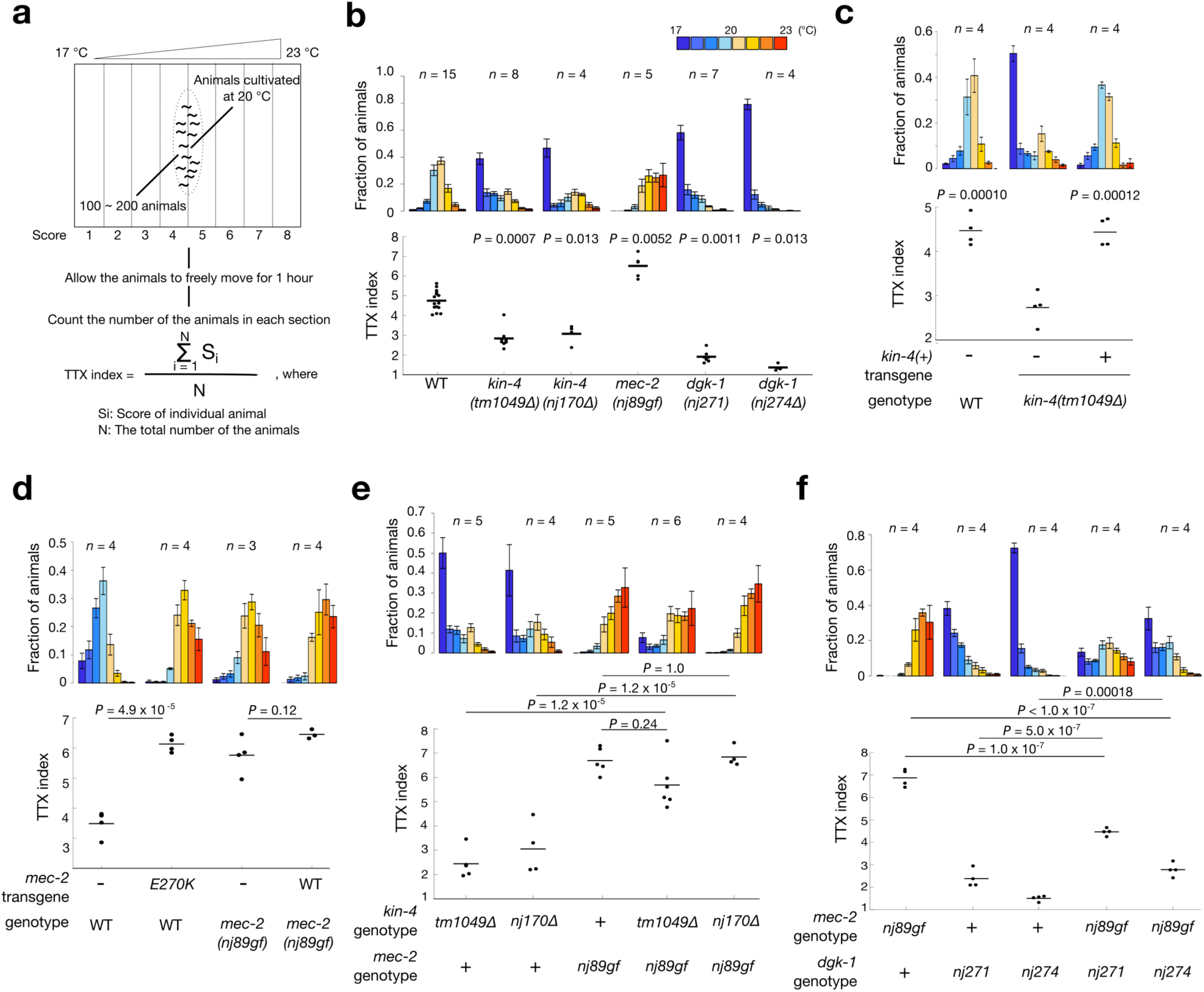
*kin-4, mec-2* and *dgk-1* Regulate the *C. elegans* Thermotaxis Behavior. (**a**) Procedure of thermotaxis assay is shown. Animals were cultivated at 20 °C and were placed on the center of a thermal gradient ranging from 17 °C to 23 °C. Each assay typically contains 100 ~ 200 animals. Distribution of the animals in each section of the assay plate was determined. We also use thermotaxis (TTX) index to quantify the behavior. The formula of TTX index is shown. (**b-f**) Thermotaxis behavior of the wild type, *kin-4*, *mec-2* and *dgk-1* mutant animals. Distributions of the animals in each section of the assay plate (top) are shown as means ± s.e.m. TTX indices (bottom) are shown as dots. Lines indicate the means. *n* represents the number of independent experiments. *P* values were determined by Kruskal-Wallis test with Steel method for comparison to the wild-type animals in (b), one-way ANOVA with Dunnett’s test for comparison to *kin-4(tm1049Δ)* mutants in (c), two-tailed Student’s *t-*tests in (d) and one-way ANOVA with Tukey-Kramer test in (e) and (f).

To address genetic interactions among these genes, we analyzed thermotaxis behaviors of double mutants. Animals carrying mutations in both *kin-4* and *mec-2* showed a thermophilic phenotype similar to that of *mec-2* single mutants (Fig. 1e), suggesting that *mec-2* acts downstream of or in parallel to *kin-4*. Similarly, *dgk-1* mutations partially suppressed the thermophilic phenotype conferred by *mec-2(nj89gf)* mutation (Fig. 1f), suggesting that *dgk-1* acts downstream of or in parallel to *mec-2*. We hereafter focused on *kin-4(tm1049Δ), mec-2(nj89gf)* and *dgk-1(nj274Δ)* mutants for further analyses.

### *kin-4, mec-2* and *dgk-1* Function in the AFD Thermosensory Neuron to Regulate Thermotaxis

To ask where *kin-4* and *mec-2* are expressed, we conducted expression analysis. We generated a functional *kin-4::gfp* translational transgene capable of rescuing the cryophilic phenotype of *kin-4* mutants (Supplementary Fig. 2a). This transgene was broadly expressed in the nervous system, and its expression was observed in neurons previously shown to be involved in thermotaxis^16,17,27^, including the AFD and AWC thermosensory neurons and the AIY and RIA interneurons (Figs. 2a and b, and Supplementary Fig. 2b). We also assessed expression of *mec-2* and found that a *Pmec-2c::gfp* reporter was expressed in AFD and AWC (Fig. 2b). As previously reported^23^, expression in mechanosensory neurons was also observed when *gfp* was fused to a promoter for another *mec-2* isoform, *mec-2a* (Supplementary Fig. 2c). A previous study also showed that *dgk-1* was ubiquitously expressed in the nervous system^25^. These results suggest that *kin-4, mec-2* and *dgk-1* are expressed in neurons known to be involved in regulation of thermotaxis, including the AFD and AWC thermosensory neurons.

**Figure 2.**
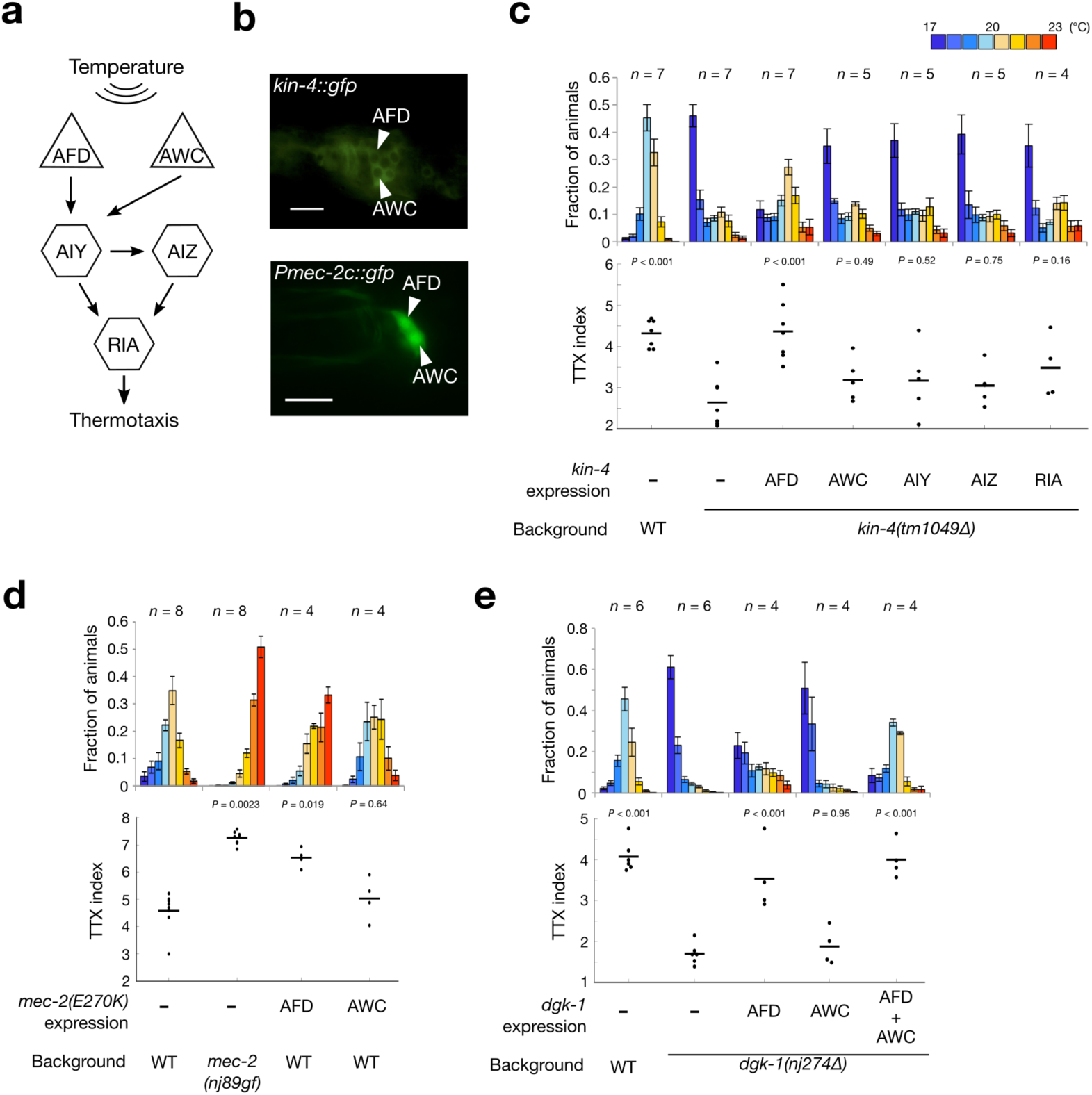
*kin-4, mec-2* and *dgk-1* Function in the AFD Thermosensory Neurons to Regulate Thermotaxis. (**a**) Neural circuit involved in thermotaxis. Arrows indicate chemical synapses. Triangles and hexagons represent sensory and interneurons, respectively. (**b**) Expression analyses of *kin-4::gfp* (Top) and *Pmec-2c::gfp* (Bottom). Head regions of animals are shown. The arrowheads indicate the AFD and AWC sensory neurons. Scale bars, 10 μm. (**c**-**e**) Thermotaxis behaviors of animals in which *kin-4, mec-2(E270K)* or *dgk-1* cDNA was specifically expressed in AFD and other neurons. Animals were cultivated at 20 °C. Distributions of animals in each section of the assay plates are shown as means ± s.e.m. TTX indices are shown as dots, and the lines indicate means. *n* indicates the number of the independent experiments. Animals were cultivated at 20 °C. *P* values were determined by one-way ANOVA with Dunnett’s test for multiple comparison to *kin-4(tm1049Δ)* in (c) or *dgk-1(nj274Δ)* in (e). Kruskal-Wallis test with Steel method was used for comparison to the wild-type animals in (d).

To identify the neuron(s) in which *kin-4, mec-2* and *dgk-1* act to regulate thermotaxis, we attempted to express each of these genes in single neurons and assessed their effects on the thermotaxis behavior. Expression of a *kin-4* cDNA in AFD rescued the cryophilic phenotype of *kin-4* mutants, whereas expression in AWC, AIY, AIZ or RIA did not (Fig. 2c), indicating that *kin-4* functions in AFD to regulate thermotaxis. When mutant *mec-2(E270K)* was expressed in AFD, it phenocopied *mec-2(nj89gf)* mutants, while expression in AWC did not (Fig. 2d). Expression of a *dgk-1* cDNA in AFD but not in AWC partially rescued the cryophilic phenotype of *dgk-1* mutants (Fig. 2e). We also observed that simultaneous expression of *dgk-1* in both AFD and AWC fully rescued the *dgk-1* mutant phenotype. These results indicate that *kin-4, mec-2* and *dgk-1* function in AFD to regulate thermotaxis.

### *kin-4, mec-2* and *dgk-1* Act Downstream of Calcium Influx in AFD

To assess whether *kin-4, mec-2* and *dgk-1* regulate temperature-evoked activity of the AFD thermosensory neuron, we conducted calcium imaging of AFD. We immobilized animals expressing the GCaMP3 calcium indicator in AFD and subjected the animals to a warming stimulus. As previously reported^20,21^, the AFD neuron showed increases in calcium concentration upon warming stimuli both below and above the cultivation temperature (Figs. 3a and b), and this response required three guanylate cyclase genes, *gcy-8, gcy-18* and *gcy-23* exclusively expressed in the AFD neurons^28–30^ (Fig. 3b). The temperature-evoked calcium responses of AFD in *kin-4, mec-2* and *dgk-1* mutants were almost indistinguishable from that of the wild-type animals (Fig. 3b-d). These results suggest that *kin-4, mec-2* and *dgk-1* regulate a process downstream of the calcium influx in AFD.

**Figure 3.**
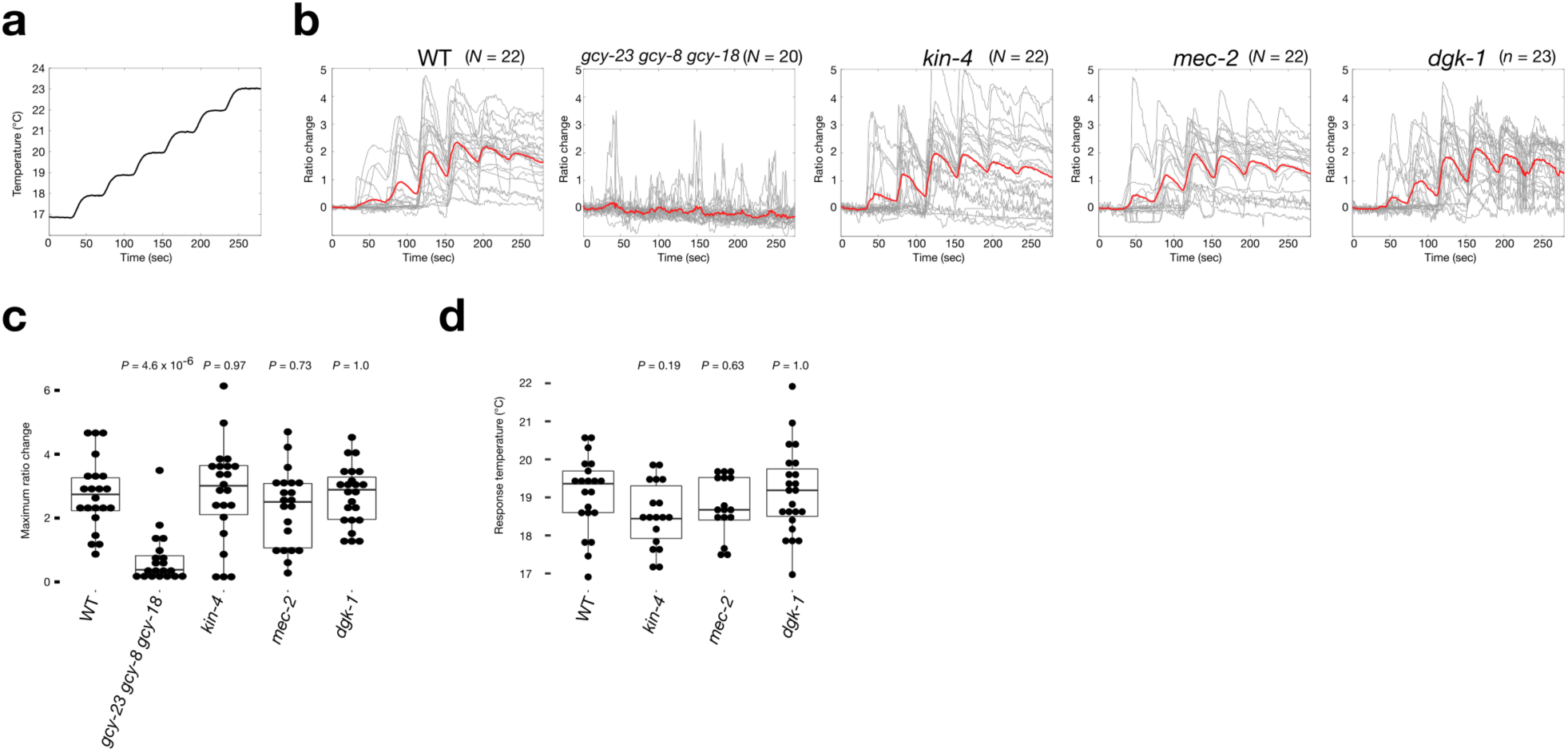
*kin-4, mec-2* and *dgk-1* Regulate a Process Downstream of Calcium Influx in AFD. (**a**) Temperature program used for calcium imaging of the AFD neurons in immobilized animals. (**b**) Ratio changes of the AFD calcium dynamics are shown. Grey lines are results for individual traces; thick colored lines represent mean values. *N* indicates the number of animals observed. (**c**) Comparison of the maximum ratio changes in each strain. Individual data points are shown as dots. Boxes display the first and third quartiles; lines inside the boxes are the medians; the whiskers extend to 1.5-time interquartile range from the box. Kruskal-Wallis test with Steel method was used to compare to the wild type animals. (**d**) Comparison of the response temperatures in the wild type, *kin-4, mec-2* and *dgk-1* mutants. The response temperature was defined previously ^68^ as the temperature at which the ratio change first exceeded 1. Individual data points are shown as dots. Boxes display the first and third quartiles; lines inside the boxes are the medians; the whiskers extend to 1.5-time interquartile range from the box. *P* values were determined by one-way ANOVA with Dunnett’s test for comparison to the wild type.

### Valence of Thermal Information Is Encoded by Bidirectional AIY Response

To ask whether *kin-4, mec-2* and *dgk-1* regulate the activity of the AIY interneuron, the sole chemical postsynaptic partner of AFD^14^, we monitored calcium dynamics of both AFD and AIY. We generated the animals expressing a calcium indicator in both AFD and AIY and subjected them to warming stimuli while they were freely moving under the microscope with an automated tracking system (Fig. 4). We first subjected animals to a warming stimulus below the cultivation temperature, in which the temperature was increasing toward the cultivation temperature (Fig. 4a). When the wild-type animals were exposed to this warming stimulus, the AFD neuron showed an increase in calcium concentration, and the AIY neuron also displayed a rise in calcium concentration after a slight drop observed at the beginning of the warming stimulus (Fig. 4b). By contrast, when the wild-type animals were subjected to a warming stimulus above the cultivation temperature, where the temperature was increasing away from the cultivation temperature (Fig. 4e), the AIY neuron instead showed a decrease in calcium concentration, despite the increase of calcium level in AFD (Fig. 4f). These results indicate that the bidirectional response of the AIY neuron correlates with the valence of thermal stimuli, with temperature increase toward the cultivation temperature evoking excitatory response and temperature increase away from the cultivation temperature inhibitory response.

**Figure 4.**
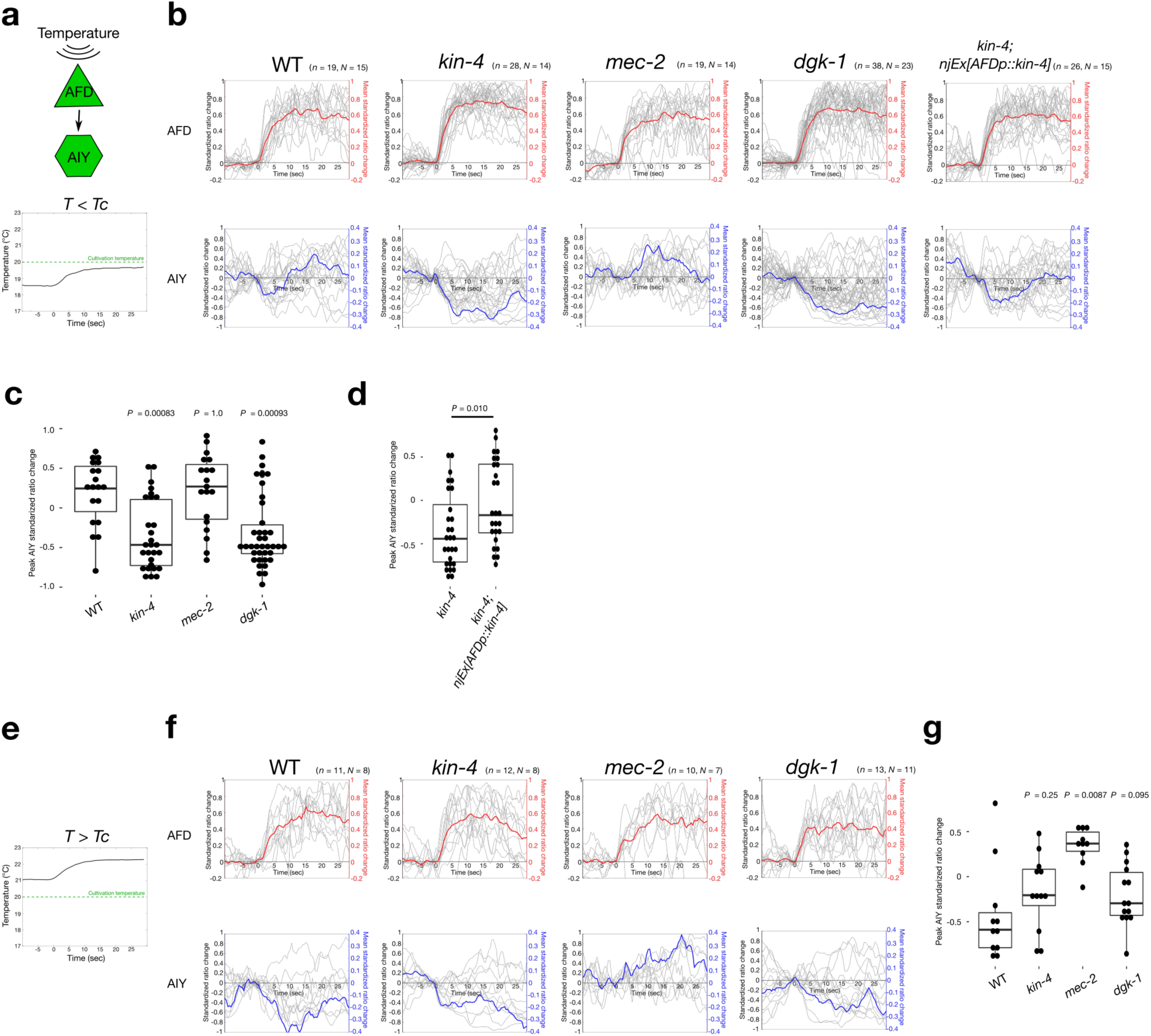
The Bidirectional AIY Response Encodes Stimulus Valence and Is Regulated by *kin-4, mec-2* and *dgk-1*. Calcium imaging of AFD and AIY neurons in freely moving animals. (**a**-**d**) Calcium imaging below the cultivation temperature. (**e**-**g**) Calcium imaging above the cultivation temperature. (**a**, **e**) Neurons imaged and the temperature program used. (**b**, **f**) Standardized ratio changes of AFD (top) and AIY (bottom) calcium dynamics. Grey lines are results for individual traces; colored lines represent mean values. Time 0 corresponds to the onset of the warming stimulus. *n* indicates the number of trials. *N* indicates the number of animals observed. Note that the scales for the mean values are on the right axis. (**c**, **d**, **g**) Comparisons of peak AIY standardized ratio changes. Individual data points are shown as dots. Boxes display the first and third quartiles; lines inside the boxes are the medians; the whiskers extend to 1.5-time interquartile range from the box. *P* values were determined by Kruskal-Wallis test with Steel method to compare to the wild type animals in (c) and (g) or by Wilcoxon rank sum test in (d).

### *kin-4, mec-2* and *dgk-1* Regulate Bidirectional AIY Response

We next examined the AIY responses in *kin-4, dgk-1* and *mec-2* mutants. The AIY neuron in *kin-4* mutants exhibited a decrease in calcium concentration even below the cultivation temperature, the condition in which the wild-type AIY neuron would normally increase its calcium level (Figs. 4b and c). Consistent with the calcium imaging analysis of AFD in immobilized animals (Fig. 3), the AFD neuron of *kin-4* mutants showed increases in calcium concentration both below and above the cultivation temperature (Figs. 4b and c). The defect in the AIY response of *kin-4* mutants was partially rescued by expression of a *kin-4* cDNA solely in the AFD neuron (Figs. 4b and d), indicating that the AIY calcium response is indeed modulated by it presynaptic partner AFD. A similar response profile was also observed in the cryophilic *dgk-1* mutants: the AIY calcium level dropped upon warming stimuli both below and above the cultivation temperature, while the AFD neuron responded to the warming stimuli by increasing the calcium concentration (Figs. 4b, c, f and g). Furthermore, the thermophilic *mec-2* mutants showed an abnormal increase in the AIY calcium concentration even above the cultivation temperature (Figs. 4f and g). Similarly in the wild type, the AFD neurons in *mec-2* mutants showed increases in calcium concentration under both conditions tested (Figs. 4b and f). These results indicate that the valence-coding bidirectional AIY activity is regulated by *kin-4, mec-2* and *dgk-1*. Given that *kin-4* expression only in AFD restored the defect of the AIY response in *kin-4* mutants, these results further suggest that presynaptic regulation is important for determining the bidirectional responses of the AIY neuron.

A recent study reported that the difference in the AIY calcium responses below or above the cultivation temperature was represented by a difference in the fraction of animals in which AIY showed an increase in calcium concentration upon warming stimulus under the experimental condition where the animals were immobilized^31^. Our results further suggest that in freely behaving animals, the AFD-AIY transmission can alter between excitatory and inhibitory communication below or above the cultivation temperature.

### Alteration of the AFD-AIY Synaptic Transmission Requires Components Essential for Neuropeptide and Glutamate Release

We next asked how the presynaptic regulation in AFD evokes opposing neuronal responses in the AIY postsynaptic neuron. Previous studies indicated that AFD employs two kinds of signaling molecules to communicate with AIY^27,32^: one is neuropeptide, and the other is glutamate. The peptidergic signaling was shown to be excitatory^32^, whereas the glutamatergic input is inhibitory due to a glutamate-gated chloride channel acting on AIY^27^. We hypothesized that the balance of these two opposing signals released from AFD might be modulated, thereby inducing an excitatory or inhibitory postsynaptic response. To test this possibility, we monitored the AIY calcium response in mutants for the gene *unc-31*, which encodes the *C. elegans* ortholog of calcium-dependent activator protein required for secretion of neuropeptides^33^. The AIY neurons of *unc-31* mutants failed to increase the calcium level and instead showed a decrease even below the cultivation temperature, while the AFD neuron showed increases in calcium concentration under both conditions tested (Figs. 5a-c). This defect in the bidirectional AIY response was partially rescued by expression of an *unc-31* cDNA only in AFD (Figs. 5b and d), indicating that the peptidergic signals from AFD is required for the bidirectional AIY response.

**Figure 5.**
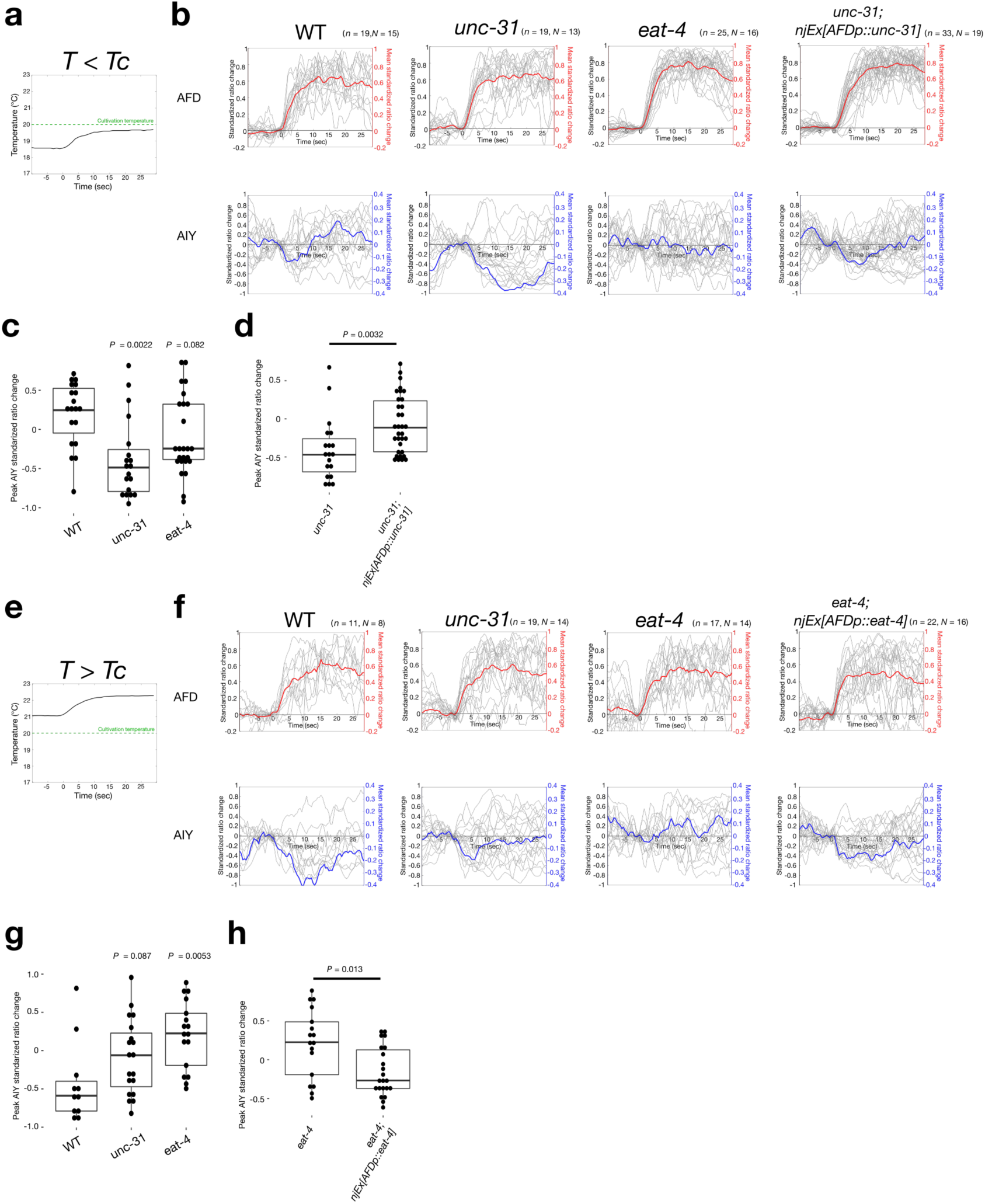
Alteration of the AFD-AIY Synaptic Valence Requires Components Essential for Neuropeptide and Glutamate Release. (**a**-**d**) Calcium imaging below the cultivation temperature. (**e**-**h**) Calcium imaging above the cultivation temperature. (**a**, **e**) The temperature program used. (**b**, **f**) Standardized ratio changes of AFD (top) and AIY (bottom) calcium dynamics. Grey lines are results for individual traces; colored lines represent mean values. Time 0 corresponds to the onset of the warming stimulus. *n* indicates the number of trials. *N* indicates the number of animals observed. Note that the scales for the mean values are on the right axis. (**c**, **d**, **g**, **h**) Comparisons of peak AIY standardized ratio changes. Individual data points are shown as dots. Boxes display the first and third quartiles; lines inside the boxes are the medians; the whiskers extend to 1.5-time interquartile range from the box. *P* values were determined by Kruskal-Wallis test with Steel method was used to compare to the wild type animals in (c) and (g) or by Wilcoxon rank sum test in (d) and (h). Note that the wild-type data are identical to those shown in Fig. 4 and indicated here for comparison to *unc-31* and *eat-4* mutants.

We also examined the AIY response in mutants for the *eat-4* gene, which encodes a *C. elegans* homolog of vesicular glutamate transporter required for transporting glutamate into synaptic vesicles^34^. In *eat-4* mutants, the AIY neurons displayed abnormal increase in the calcium concentration above the cultivation temperature (Figs. 5e-g), and this defect was rescued by expression of *eat-4* only in the AFD neurons (Figs. 5f and h). Like in *unc-31* mutants, the AFD neurons in *eat-4* mutants showed increase in the calcium level under both conditions tested. These results suggest that alteration of the positive and negative modes of the AFD-AIY communication is mediated by the presynaptic control of the glutamatergic and peptidergic outputs.

### *kin-4, mec-2* and *dgk-1* Regulate Curving Bias During Thermotaxis

To address how the alteration in the modes of the AFD-AIY communication contributes to the regulation of thermotaxis behavior, we undertook multi-worm tracking analysis^35^. We classified the animal behavior into several behavioral components discernable during *C. elegans* locomotion, such as forward locomotion, turns and reversals^36^. We sought the behavioral components that were oppositely biased below or above the cultivation temperature in the wild-type animals, and were oppositely affected by the cryophilic (*kin-4* and *dgk-1*) or thermophilic (*mec-2*) mutations. Among the behavioral components we have examined (Fig. 6), we found that the curve, change in the moving direction during forward locomotion, was one such component (Fig. 6a): the wild-type animals behaving below the cultivation temperature showed curving bias toward warmer temperature when migrating up the thermal gradient, while they curve toward colder temperature above the cultivation temperature^36^ (Fig. 6a), suggesting that this regulation of the curve would drive the animals toward the cultivation temperature. By contrast, the curving rates of the two cryophilic mutants, *kin-4* and *dgk-1*, below the cultivation temperature were abnormally biased toward the colder side (Fig. 6a). Furthermore, *mec-2* mutants behaving above the cultivation temperature failed to show a curving bias toward colder temperature (Fig. 6a). These results indicate that *kin-4, mec-2* and *dgk-1* regulate the curve during thermotaxis and suggest that the alteration of the AFD-AIY synaptic transmission generates opposing curving biases that drive the animals toward the cultivation temperature.

**Figure 6.**
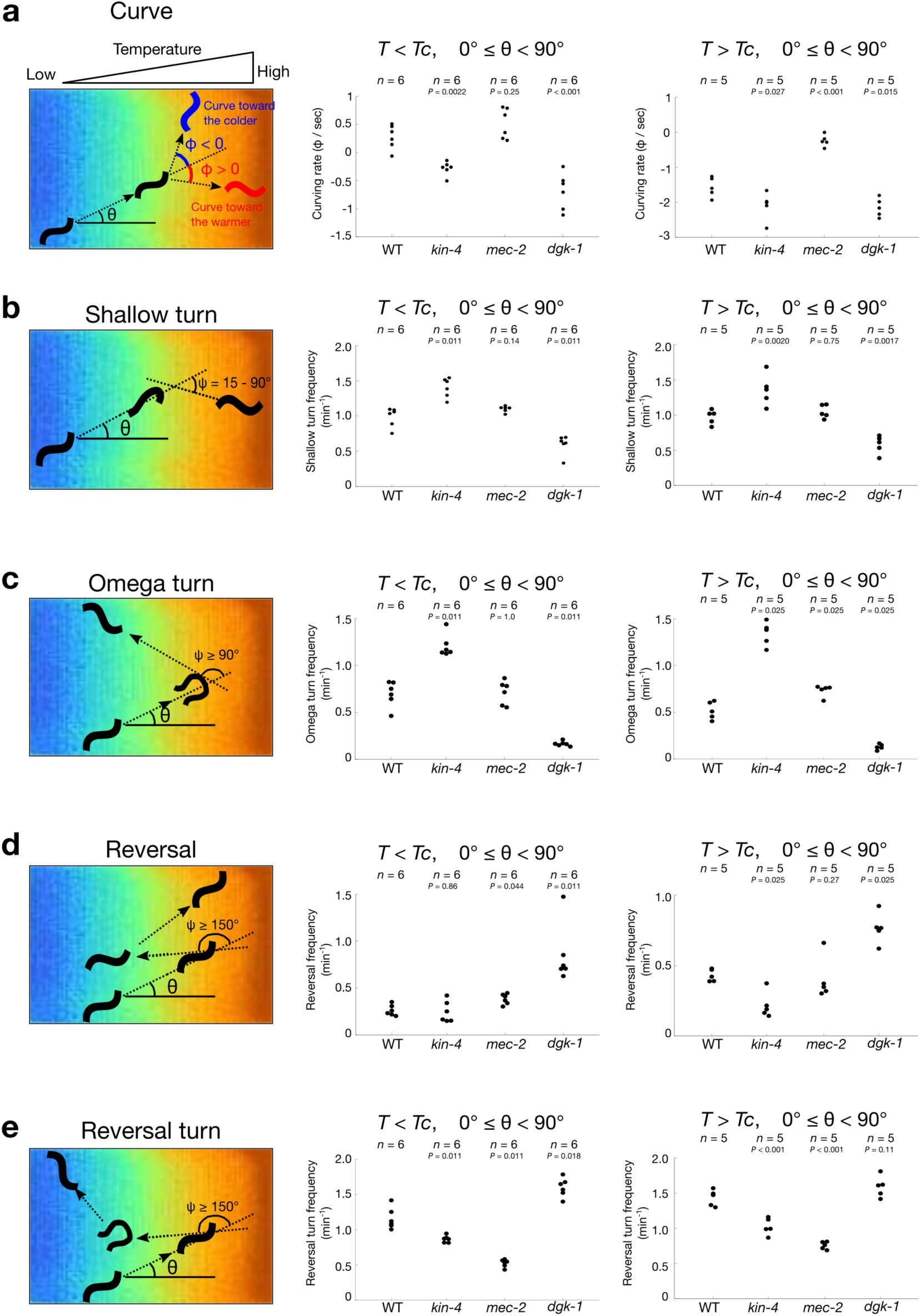
*kin-4, mec-2* and *dgk-1* Regulate Curving Bias During Thermotaxis. Multi-worm tracking analyses of curve (**a**), shallow turn (**b**), omega turn (**c**), reversal (**d**) and reversal turn (**e**). Schematics of the behavioral components are shown (left). Animals were cultivated at 20 °C (*Tc*), and their behaviors within the temperature ranges from 17.0 °C to 20.0 °C (middle) or from 20 °C to 23.0 °C (right) were monitored. (**a**) Curve is characterized by two angles ϕ and θ, where ϕ corresponds to the change in the moving direction during forward locomotion, and θ animal’s previous moving direction relative to the vector pointing to the warm side of the temperature gradient. ϕ is given as a positive value if the angle change is directed toward the warmer side and a negative value if directed toward the colder side (left). Dot plots of mean curving rate of animals migrating up the temperature gradient below (middle) or above (right) the cultivation temperature. The curving rates of all animals during the first 10 minutes of thermotaxis assay were averaged and shown as dots. *n* indicates the number of independent experiments. *P* values were determined by one-way ANOVA with Dunnett’s test for comparison to the wild-type animals. (**b**-**e**) Turns were classified into shallow turn (b) and omega turn (c), and reversals were divided into reversal (d) and reversal turn (e). The angle ψ, which represents the change in the direction after the turns or reversals, was used to classify the turns and reversals (left). Dot plots of frequencies of each behavioral component while animals were migrating up the temperature gradient below (middle) or above (right) the cultivation temperature. The frequencies of the behavioral component of all animals during the first 10 minutes of thermotaxis assay were averaged and shown as dots. *n* indicates the number of independent experiments. *P* values were determined by one-way ANOVA with Dunnett’s test or by Kruskal-Wallis test with Steel method for comparison to the wild-type animals.

## Discussion

When animals encounter environmental stimuli, they need to quickly assess whether the stimuli are beneficial or detrimental. How the brain determines whether the valence of sensory information is attractive or aversive has been a fundamental question in neurobiology. A number of previous studies indicated that the opposing valences of sensory stimuli are encoded by two separate populations of neurons, each of which represents either positive or negative valence of the stimuli^37,38^. By contrast, recent studies proposed an alternative strategy, wherein a single population of neurons responds to appetitive or aversive stimuli and represents the positive or negative valence of the stimuli by increasing or decreasing neuronal activity^39,40^. Specifically, CRF (corticotropin-releasing factor)-releasing neurons in the paraventricular nucleus of the hypothalamus are activated by aversive stimuli and inhibited by appetitive stimuli^40^. Likewise, in *C. elegans*, experience-dependent modulation enables a single set of interneurons to elicit bidirectional responses to carbon dioxide, which can be either attractive or aversive, depending on prior experience^39^. Thus, these observations indicate a previously unrecognized mechanism of valence coding for even a single modality of stimulus, in which the bidirectional activity in a single population of neurons can be modulated by prior experience and environment to represent stimulus valence. However, little was known about the molecular mechanism and circuit logics for such modulation of valence-coding activity. In this study, we report the molecular components important for determining a valence-coding activity and show that MAST kinase, Stomatin and Diacylglycerol kinase control the activity of the AIY neuron that correlates with the valence of thermal stimuli. Our results also reveal a circuit principle of such valence coding, in which a presynaptic neuron modulates its neuronal outputs and evokes an excitatory or inhibitory postsynaptic response.

We showed that *kin-4, mec-2* and *dgk-1* regulate the curving bias during thermotaxis behavior (Fig. 6). A previous study also showed that optogenetic manipulations of the AIY activity evoked biases in the curve: stimulation of AIY caused the animals to turn in the direction in which the head of the animal was bent at the time of the AIY excitation, while inhibition of AIY induced the animals to turn in the opposite direction^41^. Given this observation, our results suggest that below the cultivation temperature, a warming stimulus activates both AFD and AIY, leading to curve toward the warmer side of the temperature gradient, while above the cultivation temperature, a warming stimulus activates AFD, which in turn inhibits AIY, resulting in curving toward the colder side (Supplementary Fig. 3). Thus, the alteration of the AFD-AIY transmission mode would generate the curving bias that drives the animals toward the cultivation temperature.

Recent investigation of the global brain dynamics in *C. elegans* revealed that most interneuron layers represent the motor states of the animals^42^. Indeed, representation of the motor states is pervasive even in the first-layer interneurons that receive direct inputs from sensory neurons. The activity of the AIB interneuron, which is directly innervated by gustatory sensory neurons ASE and olfactory neurons AWC, is correlated with reversals^43^, and the AIY neuron can represent multiple motor states^41,43,44^. Because calcium dynamics within the AFD, ASE and AWC neurons represent the respective sensory stimuli^20,21,45–47^, these observations suggest that transformation of sensory information into motor representation occurs early in the neural circuit and underlies between the sensory neurons and the first-layer interneurons.

Our results indicate that the initial step of information processing that transforms thermal information into stimulus valence occurs within the AFD sensory neuron and suggest that information processing for this transformation resides in cellular processes between calcium influx and neurotransmitter release. These observations underscore neural computation at axonal regions in a single neuron. In addition to the current view that neural computations take place at the synaptic communication and dendritic computations, our results highlight the importance of axonal computation in information processing in the nervous system. These observations, in turn, raise a future challenge in neuroscience. Although electrophysiological and calcium imaging analyses undoubtedly reveal certain aspects of neuronal properties, understanding the axonal computation requires the development of methods to quantitatively measure the release of the signaling molecules, including small neurotransmitters, neuropeptides and biogenic amines^48,49^, preferably in behaving animals with high spatiotemporal resolution. Utilization of a pH-sensitive green fluorescent protein, pHluorin^50^ has been successfully used to monitor the release and recycle of synaptic vesicles^51–54^. However, since synaptic vesicle neurotransmitter content can be dynamically regulated in both vertebrates and invertebrates^55,56^, direct measurements of signaling molecules rather than that of vesicle release are required to fully understand the neuronal output. Development of such methods would unveil the dynamics of axonal computations and help dissect the mechanisms by which neurons execute this type of neural computation.

Neurons expressing multiple neurotransmitters are present in virtually all animal species. Co-transmission of multiple transmitters from single neurons can influence the same target neurons (convergent actions) or different targets (divergent actions), thereby providing additional flexibility to the circuit functions^57,58^. A few studies reported co-transmission of multiple transmitters with opposing actions^59–62^. In mammals, orexin and dynorphin neuropeptides exert opposing actions on excitability of ventral tegmental area dopamine neurons^59,60^, and in *Aplysia* neuromuscular junction, multiple co-transmitted peptides exert antagonistic actions, with one peptide promoting muscle contraction and the other increasing relaxation rate^61,62^. In these cases, multiple transmitters are co-packaged and co-released in a fixed ratio. The apparently antagonistic actions of multiple transmitters likely facilitate rhythmic muscle contraction, thereby providing temporal flexibility in the circuit function^61,62^.

Our results suggest that by altering the balance of multiple transmitters with opposing actions, a single presynaptic neuron can evoke excitatory and inhibitory postsynaptic responses, convey the valence information of sensory stimulus to the circuit, and induce an appropriate behavior. Our observations provide a sharp contrast to current views of presynaptic plasticity, in which presynaptic regulation facilitates or depresses the strength of the postsynaptic response^63,64^ but does not change the mode of transmission from an excitatory (inhibitory) to an inhibitory (excitatory) communication. We speculate that such mechanism of presynaptic control might function in other systems^65^ and be particularly effective in assigning the stimulus valence over a range of stimulus intensity. Since previous studies suggested that *dgk-1* regulates synaptic transmission at neuromuscular junction in *C. elegans*^24,25,66^*, kin-4, mec-2* and *dgk-1* might also regulate the release of glutamate and/or neuropeptide from AFD to control the bidirectional AIY response. Thus, presynaptic control of multi-transmitter release could be a fundamental mechanism of generating plasticity without extensive structural modification on the defined neural circuitry, thereby extending the computational repertoire employed by the nervous system.

## Methods

### C. elegans strains

All *C. elegans* strains were cultivated at 20 °C on nematode growth medium plates seeded with *E. coli* OP50 bacteria^67^. N2 (Bristol) was the wild-type strain. The following mutations, extrachromosomal arrays and integrated transgenes were generated or described previously. LGI: *njIs24[gcy-8p::GCaMP3, gcy-8::TagRFP]*^68^. LG III: *eat-4(ky5)*^34^. LG IV: *kin-4(tm1049Δ), kin-4(nj170Δ), unc-31(e928)*^67,69^. LG V: *njIs110[gcy-8p::4xNLS::YCX1.6, AIYp::YCX1.6]*. LG X: *mec-2(nj89gf), mec-2(nj251Δ), dgk-1(nj271), dgk-1(nj274Δ)*. Extrachromosomal arrays: *njEx682[kin-4(+), ges-1p::gfp], njEx683[kin-4::gfp, ges-1p::TagRFP], njEx753[gcy-8p::kin-4d cDNA, ges-1p::gfp*] (used for rescuing the cryophilic phenotype of *kin-4(tm1049Δ), njEx759[ttx-3p::kin-4d cDNA, ges-1p::gfp], njEx760[lin-11p::kin-4d cDNA, ges-1p::gfp], njEx763[ceh-36p::kin-4d cDNA, ges-1p::gfp], njEx764[glr-3p::kin-4d cDNA, ges-1p::gfp], njEx1029[mec-2(E270K), ges-1p::gfp], njEx1035[mec-2(+), ges-1p::gfp], njEx1158[Pmec-2a::gfp, ges-1p::TagRFP], njEx1159[Pmec-2c::gfp, ges-1p::TagRFP], njEx1220[ceh-36p::mec-2a cDNA (E270K), ceh-36p::mec-2c cDNA (E195K), ges-1p::gfp], njEx1231[gcy-8p::mec-2a cDNA (E270K), gcy-8p::mec-2c cDNA (E195K), ges-1p::gfp], njEx1317[gcy-8p::dgk-1b cDNA, ges-1p::gfp], njEx1319[odr-1p::dgk-1b cDNA, ges-1p::gfp], njEx1330[dgk-1(+), ges-1p::gfp], njEx1331[gcy-8p::dgk-1b cDNA, odr-1::dgk-1b cDNA, ges-1p::gfp], njEx1435[gcy-8p::kin-4d, ges-1p::TagRFP*] (used for rescuing AIY calcium response)*, njEx1438[gcy-8p::unc-31 cDNA, ges-1p::TagRFP], njEx1439[gcy-8p::eat-4 cDNA, ges-1p::TagRFP*]. We used following promoters for cell-specific expression of transgenes and cDNAs: *gcy-8* promoter for AFD; *ttx-3* promoter for AIY; *ceh-36* or *odr-1* promoters for AWC; *lin-11* promoter for AIZ; *glr-3* promoter for RIA; *ges-1* promoter for intestinal cells.

### Thermotaxis behavioral tests

Thermotaxis assays were performed as previously described^68^. Animals that had been cultivated at 20 °C were placed on the center of thermotaxis assay plate with a temperature gradient ranging from 17 °C to 23 °C. The steepness of the temperature gradient was set at 0.45 ~ 0.5 °C / cm. Animals were allowed to freely move on the plate for 1 hour. The thermotaxis assay plate was divided into 8 sections along the temperature gradient, and the number of adult animals in each section was counted.

### Expression analysis of *kin-4* and *mec-2*

To construct *kin-4::gfp*, we modified the fosmid containing the *kin-4* locus, WRM0635dF07. We inserted a *gfp* coding sequence immediately before the stop codon of *kin-4* using bacterial recombineering. To construct *Pmec-2a::gfp* and *Pmec-2c::gfp*, a 3.5 kb and a 3.2 kb fragments upstream of the start codon of *mec-2a* and *mec-2c* were cloned into the *gfp* expression vector pPD95.75, respectively. Animals carrying *kin-4::gfp, Pmec-2a::gfp* or *Pmec-2c::gfp* were anesthetized with 50 mM sodium azide and were observed under Nomarski optics equipped with epifluorescence. Identification of the AFD, ALM, AVM, AWC, AIY, PLM, PVM and RIA neurons was conducted by observing the positions and sizes of the nuclei and the patterns of neuronal processes.

### Multi worm tracking analysis

Multi worm tracking analysis was performed as described^36^. Animals were cultivated at 20 °C and were placed on the thermal gradient with the center temperature of 18.5 °C or 21.5 °C to monitor their behaviors below or above the cultivation temperature, respectively. The behaviors during the first 10 minutes of the thermotaxis assay were captured by a CMOS camera and were analyzed by multi worm tracker to obtain the coordinates of animal’s centroids and spines^35^. The data were further analyzed by a custom-written program in MATLAB to classify the behaviors into behavioral components.

### Calcium imaging of AFD in immobilized animals

Calcium imaging of the AFD neuron was performed as described^68^. Animals expressing GCaMP3 and TagRFP in AFD were cultivated at 20 °C and placed on a 5 – 10 % agarose pad with polystyrene beads to immobilize the animals. Animals were subjected to a warming stimulus, and the fluorescence intensities from the AFD cell body were captured and analyzed by the MetaMorph software. The ratio of the fluorescence intensity (GCaMP3/TagRFP) was used to calculate ratio change, which was defined as (ratio – baseline ratio)/baseline ratio, where baseline ratio was the mean of the ratio values during the first 30 seconds of the experiment. The response temperature was previously defined^68^ as the temperature at which the ratio change first exceeded 1.

### Calcium imaging of AIY in freely moving animals

We generated animals expressing the ratiometric calcium probe YCX1.6 in AFD and AIY. YCX1.6 in AFD was localized to the nucleus to separate the fluorescence signals from these neurons. Animals were cultivated at 20 °C and placed on a 2 – 2.5 % agarose pad on a cover glass. The sample was covered by another cover glass and was placed onto a transparent temperature-controlled device (TOKAI HIT Co. Ltd., Fujinomiya). This device was installed onto a motorized stage (HawkVision Inc., Fujisawa) that keeps the target image of animals in the field of view. Controlling the stage movement was achieved by real-time analysis of transmitted infrared light images. The fluorescence images were captured twice a second and split into YFP and CFP channels by W-VIEW GEMINI (Hamamatsu photonics K.K., Hamamatsu). The YFP and CFP fluorescence intensities were obtained from the cell body of AFD and an axonal region of AIY^68^. The fluorescent images were analyzed by a custom-written program in MATLAB with manual inspection of region of interest in every frame, and the fluorescent intensities of YFP and CFP were determined. We eliminated from the analysis the trials in which the temperature program failed to activate AFD. For this purpose, we applied a low-pass filter to the YFP/CFP ratio with the cut-off frequency at 0.05 Hz, and the resulting ratio data were used to calculate the ratio change. We eliminated from the analysis the trails in which the maximum ratio changes were smaller than 0.25 for recordings below the cultivation temperature or 0.22 for above the cultivation temperature. These threshold values were determined from experiments in which the temperature was kept constant. For the reminder of the trials, the ratio of fluorescence intensities (YFP/CFP) was used to calculate the standardized ratio change of AFD and AIY, which was defined as (ratio – minimum ratio)/(maximum ratio – minimum ratio). The baseline standardized ratio, which was the mean of the standardized ratio values of 5 consecutive frames immediately before the onset of warming stimulus, was subtracted from the standardized ratio change of each frame. For comparison of the peak standardized ratio change, the maximum standardized ratio change (positive or negative) in a 2-second time window centered at the peak of the mean control standardized ratio change was used.

### Statistics

Normality of the data was assessed by Shapiro-Wilk test. Equal variance among data sets was assessed by *F*-test or Bartlett test. When both normality and equal variance were assumed for the data set, we used two-tailed student *t-*test for pairwise comparison and one-way analysis of variance (ANOVA) with Tukey-Kramer or Dunnett’s test for multiple comparisons. In other cases, we applied Wilcoxon rank sum test for pairwise comparison and Kruskal-Wallis rank sum test with Steel method for multiple comparisons.

## Data availability

All raw images, source data and custom scripts are available from the authors upon reasonable request.

## Acknowledgments

We thank Y. Kohara, K. G. Miller for cDNAs; S. Mitani at National BioResouce for strains; K. Noma for comments on this manuscript; K. Ikegami, Y. Murakami, J. Okada, T. Sakaki, K. Sawayama, F. Takeshige for technical assistance. M.I. was supported by KAKENHI 16J05770. This work was supported by JSPS KAKENHI Grant Numbers 17K07499 (to S.N.), 18H05123 (to S.N.), 26560549 (to Y.T.), 16H06536 (to K.H.) 18H04693 (to I.M.), 16H01272 (to I.M.), 16H02516 (to I.M.) and by ERATO project (JPMJER1004 to TH) from JST.

## Author Contributions

S.N. and I.M. designed the experiments. S.N., R.A., A.S. and R.K. conducted experiments. S.N. and M.I. wrote custom codes for the analyses. Y.T., X.F. and K.H. developed and set up the tracking system for calcium imaging of freely moving animals. T. S., K. I. and T. H. conducted whole genome sequencing of mutants isolated in this study. S.N. and I. M. wrote the manuscript.

## Materials & Correspondence

Correspondence and requests for materials should be addressed to I.M.

## Supplemental Information

### Supplemental Figure Legends

**Supplemental Figure 1.**
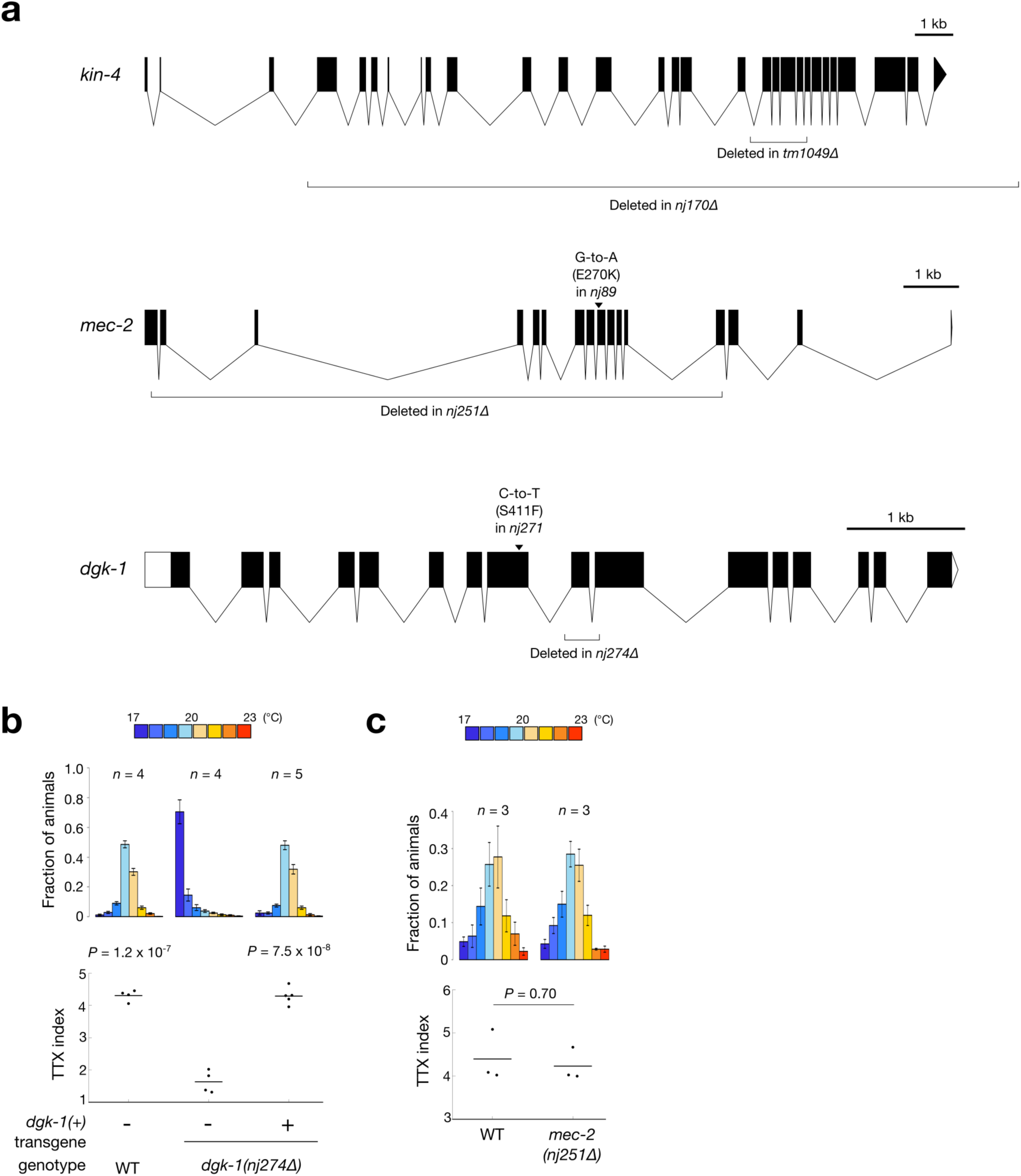
*kin-4, mec-2* and *dgk-1* Regulate Thermotaxis Behavior. (**a**) Gene structures of *kin-4* (top), *mec-2* (middle) and *dgk-1* (bottom) and mutations associated with each mutant are shown. Filled boxes and triangles represent exons, empty boxes and triangles untranslated region, and lines introns. (**b**, **c**) Thermotaxis behaviors of *dgk-1(nj274Δ)* carrying a genomic *dgk-1(+)* transgene (b) and *mec-2(nj251Δ)* mutants (c). Animals were cultivated at 20 °C and were placed on a thermal gradient ranging from 17 °C to 23 °C. Distributions of the animals (top) in each section of the assay plates are shown as means ± s.e.m. TTX indices (bottom) are shown as dots. Lines indicate the means. *P* values were determined by one-way ANOVA with Dunnett’s test for comparison to *dgk-1(nj274Δ)* mutants (b) and two-tailed Student’s *t-*tests in (c).

**Supplemental Figure 2.**
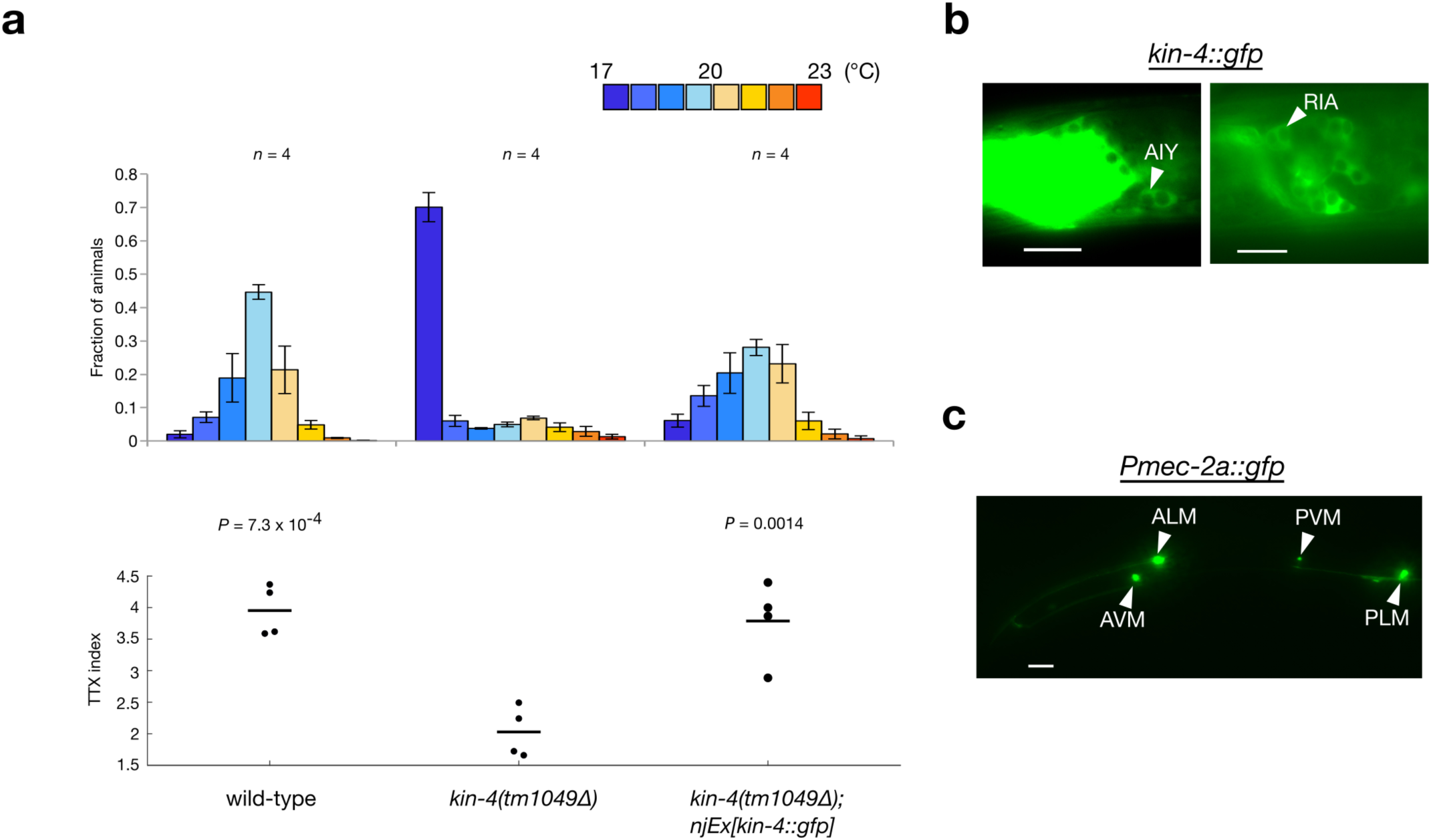
Expression analysis of *kin-4::gfp* and *Pmec-2::gfp*. (**a**) Thermotaxis behavior of animals carrying a translational *kin-4::gfp* fusion. Wild-type, *kin-4(tm1049Δ)* mutants and *kin-4(tm1049Δ)* animals carrying the *kin-4::gfp* transgene were cultivated at 20 °C and subjected to thermotaxis assay. Distributions of the animals in each section of the assay plates are shown as means ± s.e.m (top). *n* indicates the number of the independent experiments. TTX indices are shown as dots. Lines indicate the means. *P* values were determined by one-way ANOVA with Dunnett’s test for comparison to *kin-4(tm1049Δ)* animals. (**b**) Expression analysis of the *kin-4::gfp* transgene. Head regions of animals are shown. The *kin-4::gfp* fusion was expressed in the AIY (left) and RIA (right) interneurons. Scale bars, 10 μm. (**c**) Expression analysis of *Pmec-2a::gfp*. The entire body of an animal is shown. The *Pmec-2a::gfp* reporter was expressed in the ALM, AVM, PLM and PVM neurons. Scale bar, 50 μm.

**Supplemental Figure 3.**
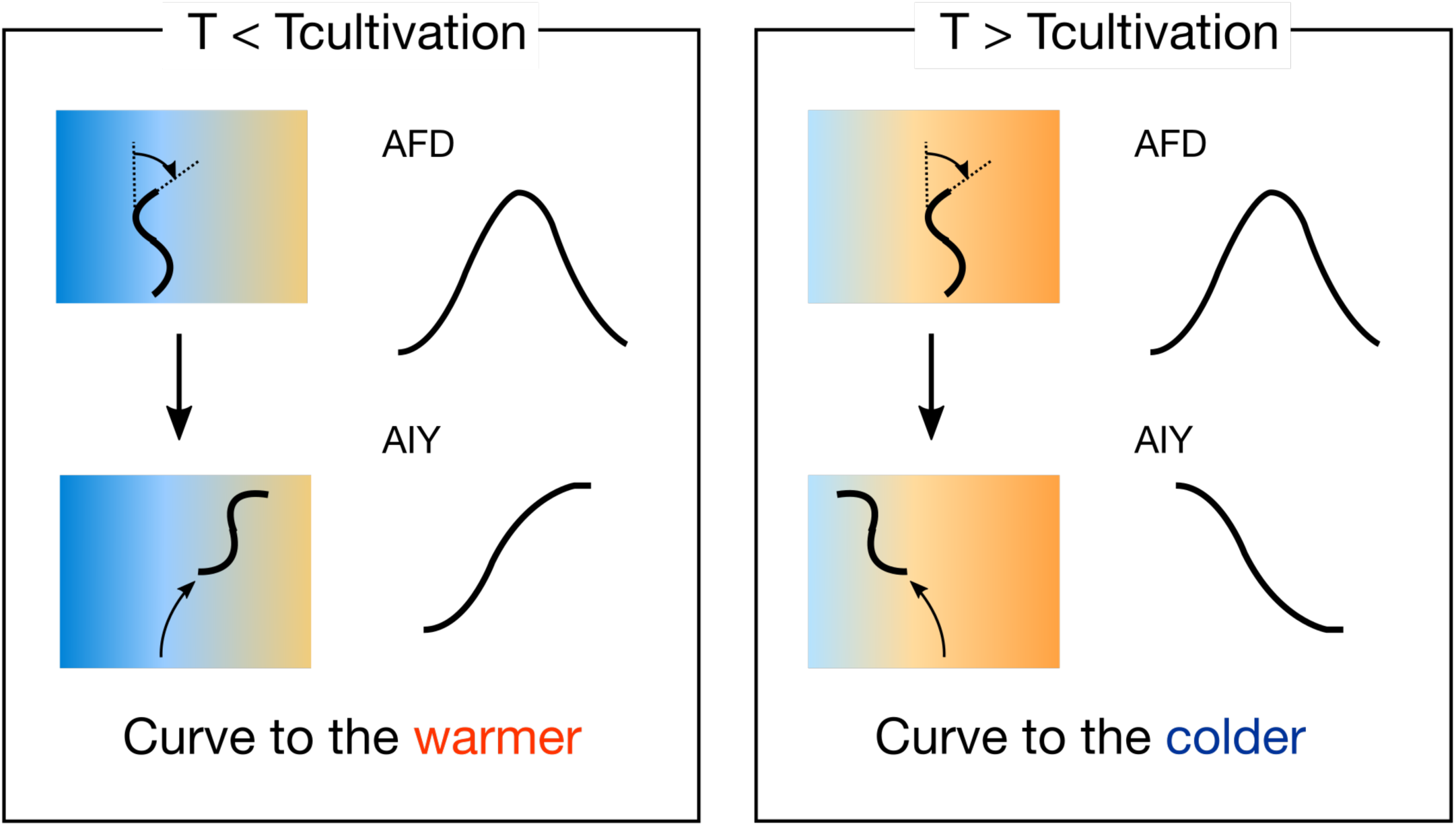
Model of curve regulation by alteration of AFD-AIY neuronal communication. Below the cultivation temperature (left), a head bending toward warmer side of the temperature gradient activates both AFD and AIY, leading to curve toward warmer side. Above the cultivation temperature (right), a temperature increase activates AFD, which in turn, inhibits AIY, leading to curve toward colder side.

## References

1. Zangrossi, H. & File, S. E. Behavioral consequences in animal tests of anxiety and exploration of exposure to cat odor. Brain Res. Bull. 29, 381–388 (1992).

2. Terry, L. M. & Johanson, I. B. Olfactory influences on the ingestive behavior of infant rats. Dev. Psychobiol. 20, 313–331 (1987).

3. Johnston, R. E. Effects of female odors on the sexual behavior of male hamsters. Behav. Neural Biol. 46, 168–188 (1986).

4. Min, S., Ai, M., Shin, S. A. & Suh, G. S. B. Dedicated olfactory neurons mediating attraction behavior to ammonia and amines in *Drosophila*. Proc. Natl. Acad. Sci. 110, E1321–E1329 (2013).

5. Kobayakawa, K. et al. Innate versus learned odour processing in the mouse olfactory bulb. Nature 450, 503–508 (2007).

6. Mueller, K. L. et al. The receptors and coding logic for bitter taste. Nature (2005). doi:10.1038/nature03352

7. Zhao, G. Q. et al. The receptors for mammalian sweet and umami taste. Cell 115, 255–266 (2003).

8. Marella, S. et al. Imaging taste responses in the fly brain reveals a functional map of taste category and behavior. Neuron 49, 285–295 (2006).

9. Laing, D. G., Panhuber, H. & Baxter, R. I. Olfactory properties of amines and n-butanol. Chem. Senses 3, 149–166 (1978).

10. Charro, M. J. & Alcorta, E. Quantifying relative importance of maxillary palp information on the olfactory behavior of *Drosophila melanogaster*. J. Comp. Physiol. A 175, 761–766 (1994).

11. Malnic, B., Hirono, J., Sato, T. & Buck, L. B. Combinatorial receptor codes for odors. Cell 96, 713–723 (1999).

12. Semmelhack, J. L. & Wang, J. W. Select *Drosophila* glomeruli mediate innate olfactory attraction and aversion. Nature 459, 218–223 (2009).

13. Yoshida, K. et al. Odour concentration-dependent olfactory preference change in *C. elegans*. Nat. Commun. 3, 711–739 (2012).

14. White, J. G., Southgate, E., Thomson, J. N. & Brenner, S. The Structure of the Nervous System of the Nematode *Caenorhabditis elegans*. Philos. Trans. R. Soc. B Biol. Sci. 314, 1–340 (1986).

15. Hedgecock, E. M. & Russell, R. L. Normal and mutant thermotaxis in the nematode *Caenorhabditis elegans*. Proc. Natl. Acad. Sci. U. S. A. 72, 4061–4065 (1975).

16. Kuhara, A. et al. Temperature sensing by an olfactory neuron in a circuit controlling behavior of *C. elegans*. Science 320, 803–807 (2008).

17. Mori, I. & Ohshima, Y. Neural regulation of thermotaxis in *Caenorhabditis elegans*. Nature 376, 344–348. (1995).

18. Beverly, M., Anbil, S. & Sengupta, P. Degeneracy and Neuromodulation among Thermosensory Neurons Contribute to Robust Thermosensory Behaviors in *Caenorhabditis elegans*. J. Neurosci. 31, 11718–11727 (2011).

19. Luo, L. et al. Bidirectional thermotaxis in *Caenorhabditis elegans* is mediated by distinct sensorimotor strategies driven by the AFD thermosensory neurons. Proc. Natl. Acad. Sci. 111, 2776–2781 (2014).

20. Kimura, K. D., Miyawaki, A., Matsumoto, K. & Mori, I. The *C. elegans* thermosensory neuron AFD responds to warming. Curr. Biol. 14, 1291–1295 (2004).

21. Clark, D. A., Biron, D., Sengupta, P. & Samuel, A. D. T. The AFD Sensory Neurons Encode Multiple Functions Underlying Thermotactic Behavior in *Caenorhabditis elegans*. J. Neurosci. 26, 7444–7451 (2006).

22. Walden, P. D. & Cowan, N. J. A Novel 205-Kilodalton Testis-Specific Serine/Threonine Protein Kinase Associated with Microtubules of the Spermatid Manchette. Mol. Cell. Biol. 13, 7625–7635 (1993).

23. Huang, M., Gu, G., Ferguson, E. L. & Chalfie, M. A stomatin-like protein necessary for mechanosensation in *C. elegans*. Nature 378, 292–295 (1995).

24. Miller, K. G., Emerson, M. D. & Rand, J. B. Goalpha and diacylglycerol kinase negatively regulate the Gqalpha pathway in *C. elegans*. Neuron 24, 323–333 (1999).

25. Nurrish, S., Ségalat, L. & Kaplan, J. M. Serotonin inhibition of synaptic transmission: gα(o) decreases the abundance of *unc-13* at release sites. Neuron 24, 231–242. (1999).

26. Okochi, Y., Kimura, K. D., Ohta, A. & Mori, I. Diverse regulation of sensory signaling by *C. elegans* nPKC-epsilon/eta TTX-4. EMBO J. 24, 2127–2137 (2005).

27. Ohnishi, N., Kuhara, A., Nakamura, F., Okochi, Y. & Mori, I. Bidirectional regulation of thermotaxis by glutamate transmissions in *Caenorhabditis elegans*. EMBO J. 30, 1376–1388 (2011).

28. Inada, H. et al. Identification of guanylyl cyclases that function in thermosensory neurons of *Caenorhabditis elegans*. Genetics 172, 2239–2252 (2006).

29. Ramot, D., MacInnis, B. L. & Goodman, M. B. Bidirectional temperature-sensing by a single thermosensory neuron in *C. elegans*. Nat. Neurosci. 11, 908–915 (2008).

30. Takeishi, A. et al. Receptor-type Guanylyl Cyclases Confer Thermosensory Responses in *C. elegans*. Neuron 90, 235–244 (2016).

31. Hawk, J. D. et al. Integration of Plasticity Mechanisms within a Single Sensory Neuron of *C. elegans* Actuates a Memory. Neuron 356–367 (2018). doi:10.1016/j.neuron.2017.12.027

32. Narayan, A., Laurent, G. & Sternberg, P. W. Transfer characteristics of a thermosensory synapse in *Caenorhabditis elegans*. Proc. Natl. Acad. Sci. 108, 9667–9672 (2011).

33. Speese, S. et al. UNC-31 (CAPS) Is Required for Dense-Core Vesicle But Not Synaptic Vesicle Exocytosis in *Caenorhabditis elegans*. J. Neurosci. 27, 6150–6162 (2007).

34. Lee, R. Y. N., Sawin, E. R., Chalfie, M., Horvitz, H. R. & Avery, L. EAT-4, a Homolog of a Mammalian Sodium-Dependent Inorganic Phosphate Cotransporter, Is Necessary for Glutamatergic Neurotransmission in *Caenorhabditis elegans*. Neuron 19, 159–167 (1999).

35. Swierczek, N. A., Giles, A. C., Rankin, C. H. & Kerr, R. A. High-throughput behavioral analysis in *C. elegans*. Nat. Methods 8, 592–598 (2011).

36. Ikeda, M. et al. Circuit Degeneracy Facilitates Robustness and Flexibility of Navigation Behavior in *C. elegans*. bioRxiv (2018). doi:10.1101/385468

37. Bruchas, M. R., Calhoon, G. G., Al-Hasani, R., Namburi, P. & Tye, K. M. Architectural Representation of Valence in the Limbic System. Neuropsychopharmacology 41, 1697–1715 (2015).

38. Knaden, M. & Hansson, B. S. Mapping odor valence in the brain of flies and mice. Curr. Opin. Neurobiol. 24, 34–38 (2014).

39. Guillermin, M. L., Carrillo, M. A. & Hallem, E. A. A Single Set of Interneurons Drives Opposite Behaviors in *C. elegans*. Curr. Biol. 27, 2630–2639.e6 (2017).

40. Kim, J. et al. Rapid, biphasic CRF neuronal responses encode positive and negative valence. Nat. Neurosci. (2019). doi:10.1038/s41593-019-0342-2

41. Kocabas, A., Shen, C. H., Guo, Z. V. & Ramanathan, S. Controlling interneuron activity in *Caenorhabditis elegans* to evoke chemotactic behaviour. Nature 490, 273–277 (2012).

42. Kato, S. et al. Global Brain Dynamics Embed the Motor Command Sequence of *Caenorhabditis elegans*. Cell 163, 656–669 (2015).

43. Luo, L. et al. Dynamic encoding of perception, memory, and movement in a *C. elegans* chemotaxis circuit. Neuron 82, 1115–1128 (2014).

44. Li, Z., Liu, J., Zheng, M. & Xu, X. Z. S. Encoding of both analog- and digital-like behavioral outputs by one *C. elegans* interneuron. Cell 159, 751–765 (2014).

45. Suzuki, H. et al. Functional asymmetry in *Caenorhabditis elegans* taste neurons and its computational role in chemotaxis. Nature 454, 114–117 (2008).

46. Chalasani, S. H. et al. Dissecting a circuit for olfactory behaviour in *Caenorhabditis elegans*. Nature 450, 63–70 (2007).

47. Kunitomo, H. et al. Concentration memory-dependent synaptic plasticity of a taste circuit regulates salt concentration chemotaxis in *Caenorhabditis elegans*. Nat. Commun. 4, 1–11 (2013).

48. Marvin, J. S. et al. An optimized fluorescent probe for visualizing glutamate neurotransmission. Nat. Methods 10, 162–170 (2013).

49. Sun, F. et al. A Genetically Encoded Fluorescent Sensor Enables Rapid and Specific Detection of Dopamine in Flies, Fish, and Mice. Cell 174, 481–496 (2018).

50. Miesenbock, G., Rothman, D. A. D. A. & Rothman, J. E. Visualizingsecretionand synaptictransmissionwith pH-sensitivegreen fluorescent proteins. Nature 394, 192–195 (1998).

51. Bozza, T., McGann, J. P., Mombaerts, P. & Wachowiak, M. In vivo imaging of neuronal activity by targeted expression of a genetically encoded probe in the mouse. Neuron 42, 9–21 (2004).

52. Li, Z. et al. Synaptic vesicle recycling studied in transgenic mice expressing synaptopHluorin. Proc. Natl. Acad. Sci. 102, 6131–6136 (2005).

53. Ventimiglia, D. & Bargmann, C. I. Diverse modes of synaptic signaling, regulation, and plasticity distinguish two classes of *C. elegans* glutamatergic neurons. Elife 6, 1–25 (2017).

54. Sankaranarayanan, S. & Ryan, T. A. Calcium accelerates endocytosis of vSNAREs at hippocampal synapses. Nat. Neurosci. 4, 129–136 (2001).

55. Aguilar, J. I. et al. Neuronal Depolarization Drives Increased Dopamine Synaptic Vesicle Loading via VGLUT. Neuron 95, 1074–1088.e7 (2017).

56. Steinert, J. R. et al. Experience-Dependent Formation and Recruitment of Large Vesicles from Reserve Pool. Neuron 50, 723–733 (2006).

57. Nusbaum, M. P., Blitz, D. M. & Marder, E. Functional consequences of neuropeptide and small-molecule co-transmission. Nat. Rev. Neurosci. 18, 389–403 (2017).

58. Vaaga, C. E., Borisovska, M. & Westbrook, G. L. Dual-transmitter neurons: Functional implications of co-release and co-transmission. Curr. Opin. Neurobiol. 29, 25–32 (2014).

59. Baimel, C., Lau, B. K., Qiao, M. & Borgland, S. L. Projection-Target-Defined Effects of Orexin and Dynorphin on VTA Dopamine Neurons. Cell Rep. 18, 1346–1355 (2017).

60. Muschamp, J. W. et al. Hypocretin (orexin) facilitates reward by attenuating the antireward effects of its cotransmitter dynorphin in ventral tegmental area. Proc. Natl. Acad. Sci. 111, E1648–E1655 (2014).

61. Vilim, F. S., Price, D. a, Lesser, W., Kupfermann, I. & Weiss, K. R. Costorage and corelease of modulatory peptide cotransmitters with partially antagonistic actions on the accessory radula closer muscle of *Aplysia californica*. J. Neurosci. 16, 8092–104 (1996).

62. Vilim, F. S., Cropper, E. C., Price, D. A., Kupfermann, I. & Weiss, K. R. Peptide Cotransmitter Release from Motorneuron B16 in *Aplysia californica*: Costorage, Corelease, and Functional Implications. J. Neurosci. 20, 2036–2042 (2000).

63. Monday, H. R. & Castillo, P. E. Closing the gap: long-term presynaptic plasticity in brain function and disease. Curr. Opin. Neurobiol. 45, 106–112 (2017).

64. Regehr, W. G. Short-term presynaptic plasticity. Cold Spring Harb. Perspect. Biol. 4, a005702 (2012).

65. Tsunozaki, M., Chalasani, S. H. & Bargmann, C. I. A Behavioral Switch: cGMP and PKC Signaling in Olfactory Neurons Reverses Odor Preference in *C. elegans*. Neuron 59, 959–971 (2008).

66. McMullan, R., Hiley, E., Morrison, P. & Nurrish, S. J. Rho is a presynaptic activator of neurotransmitter release at pre-existing synapses in *C. elegans*. Genes Dev. 20, 65–76 (2006).

67. Brenner, S. The genetics of *Caenorhabditis elegans*. Genetics 77, 71–94 (1974).

68. Kobayashi, K. et al. Single-Cell Memory Regulates a Neural Circuit for Sensory Behavior. Cell Rep. 14, 11–21 (2016).

69. Charlie, N. K., Schade, M. A., Thomure, A. M. & Miller, K. G. Presynaptic UNC-31 (CAPS) is required to activate the Gαs pathway of the *Caenorhabditis elegans* synaptic signaling network. Genetics 172, 943–961 (2006).

